# Metagenomic analysis of ecological niche overlap and community collapse in microbiome dynamics

**DOI:** 10.1101/2023.01.17.524457

**Authors:** Hiroaki Fujita, Masayuki Ushio, Kenta Suzuki, Masato S. Abe, Masato Yamamichi, Yusuke Okazaki, Alberto Canarini, Ibuki Hayashi, Keitaro Fukushima, Shinji Fukuda, E. Toby Kiers, Hirokazu Toju

## Abstract

Species utilizing the same resources often fail to coexist for extended periods of time. Such competitive exclusion mechanisms potentially underly microbiome dynamics, causing breakdowns of communities composed of species with similar genetic backgrounds of resource utilization. Although genes responsible for competitive exclusion among a small number of species have been investigated in pioneering studies, it remains a major challenge to integrate genomics and ecology for understanding stable coexistence in species-rich communities. Here, we show that community-scale analyses of functional gene redundancy can provide a useful platform for interpreting and predicting collapse of bacterial communities. Through 110-day time-series of experimental microbiome dynamics, we analyzed the metagenome-assembled genomes of co-occurring bacterial species. We then inferred ecological niche space based on the multivariate analysis of the genome compositions. The analysis allowed us to evaluate potential shifts in the level of niche overlap between species through time. We hypothesized that community-scale pressure of competitive exclusion could be evaluated by quantifying overlap of genetically determined resource-use profiles (metabolic pathway profiles) among coexisting species. We found that the degree of community compositional changes observed in the experimental microbiome was explained by the magnitude of gene-repertoire overlaps among bacterial species. The metagenome-based analysis of genetic potential for competitive exclusion will help us forecast major events in microbiome dynamics such as sudden community collapse (i.e., dysbiosis).

## INTRODUCTION

Classic niche theory predicts that coexistence of species requires interspecific difference in resource use patterns [1–6]. Although some specific mechanisms can promote stable coexistence even with complete resource overlap (e.g., spatial structure of habitats and temporal variability in resource availability), similarity/dissimilarity in basic resource dependency among species is the basic factor determining the occurrence of competitive exclusion [7–9]. Therefore, evaluating the overlap of “fundamental niches”, which are defined by species’ fundamental resource requirements and resource-use capabilities [10, 11], is an essential step for understanding and predicting community-level dynamics.

Insights into fundamental niches are encrypted in species’ genomes [12–14]: as species’ traits are encoded in their DNA, genomic information provides the ultimate basis for evaluating target species’ fundamental niches [15, 16]. Thus, potential strength of competitive interactions within ecological guilds or communities could be evaluated based on the distribution of species’ gene repertoires within ecological niche space inferred with metagenomic data [12, 15, 16], also referred to as “metagenomic niche space”. Although overlap of niches does not always cause competitive exclusion [7–9], higher levels of gene repertoire overlap within a community may impose greater impacts on population dynamics of constituent species.

In considering coexistence of microbial species, it is essential to examine whether such competition-driven population-level phenomena underly drastic ecological events observed at the community level. Microbial communities sometimes show sudden and substantial changes in species and/or taxonomic compositions [17–20]. Human gut microbiomes, for example, have been reported to show drastic shifts from species-rich states to “imbalanced” states with low *α*-diversity and overrepresentation of pathogenic species [21–24] (e.g., *Clostridium difficile*). Elucidating the ecological mechanisms causing such drastic community-level events provide fundamental insights into microbiome dynamics [24–26]. In this respect, an important challenge is to test the hypothesis that high levels of gene-repertoire overlap are observable prior to drastic community compositional changes. However, this hypothesis, to our knowledge, has not yet been tested presumably due to the paucity of time-series observations of microbiomes with substantial compositional changes. Even if such microbiome time-series data are available, analyses of potential niche (gene repertoire) overlap require another line of information. Specifically, we need data of respective species’ genomes at multiple time points. Therefore, developing research systems that can overcome these constrains will deepen our understanding of microbiome ecological processes.

In this study, we test the degree to which gene-repertoire overlap changes through dynamics of species-rich microbial communities. By targeting an experimental microbial system showing rapid and substantial changes in taxonomic compositions [19], we infer niche space depicting species’ gene repertoires. Work in this system using a metabolic modeling analysis demonstrated that interactions between species were keys to understand the drastic microbiome dynamics [27]. Now, by compiling the shotgun metagenomic data collected at 13 time points across the 110-day time-series of the experiment, we reveal temporal shifts in the magnitude of gene repertoire overlap among microbial species. We then examine whether a high level of fundamental-niche overlap is observed prior to drastic changes in community structure. Overall, we explore how signs of drastic shifts in community structure are detected by inferring community-scale degree of fundamental niche overlap with the aid of genomic information.

## RESULTS

### Functional dynamics of microbiomes

We focused on the experimental microbiome showing drastic shifts in taxonomic compositions [19]. In a previous study [19], a 110-day monitoring of microbiomes was performed with six experimental settings. To set up experimental microbiomes with high diversity of bacterial species/taxa, we used natural microbial communities derived from soil or pond-water ecosystems as source inocula, rather than “synthetic” communities with pre-defined diversity. Specifically, microbiomes were set up with combinations of two source inoculum types (soil-or pond-water-derived inoculum microbiomes) and three medium types (oatmeal, oatmeal-peptone, or peptone broth medium) with eight replications (2 × 3 × 8 = 48 microbiomes; see Materials and Methods for details). From each of the 48 microbiomes, a fraction of each replicate community was sampled every 24 hours. The collected samples were subjected to the amplicon sequencing of the 16S rRNA region and the temporal changes in community compositions were monitored throughout the time-series [19]. By calculating the magnitude of time-series changes in community compositions [19] (Fig. 1A), we focused on a water-inoculum/oatmeal-medium replicate community showing the most abrupt (rapid and substantial) changes in community compositions among the 48 microbiomes examined as described in a study on metabolic interactions between species [27] (Fig. S1).

**Fig. 1.**
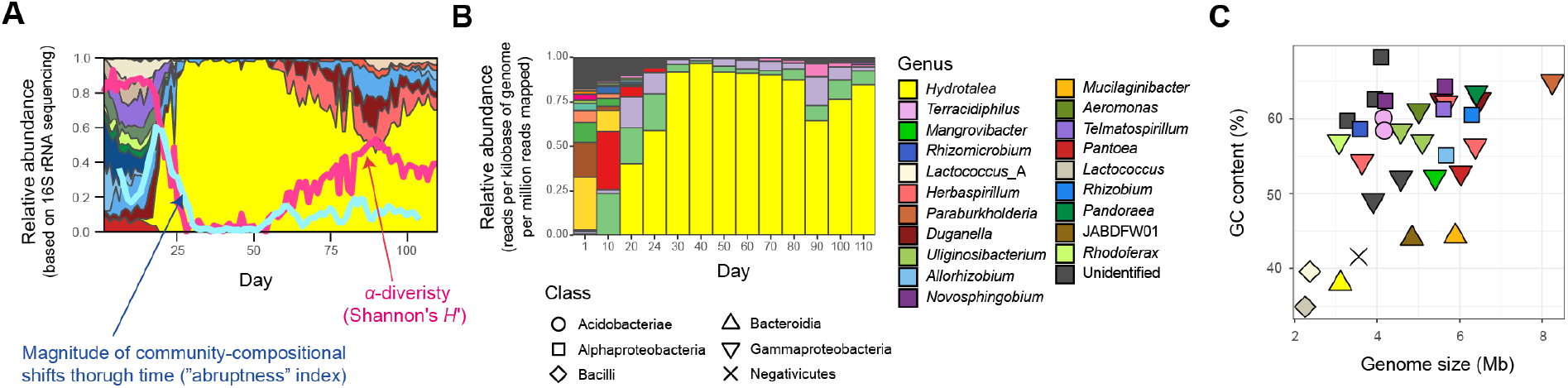
Community and ecosystem dynamics. **A** Time-series data of community structure. For the replicate microcosm that showed the most abrupt community compositional changes through the 110-day microbiome experiment [19] (Fig. S1), family-level taxonomic compositions inferred with 16S rRNA sequencing are shown. The blue line represents the speed and magnitude of community compositional changes around each time point (“abruptness” index [19]; see Materials and Methods). The red line indicates *α*-diversity (Shannon’s *H’*) of microbial ASVs [19]. Note that a value larger than 0.5 represents turnover of more than 50 % of microbial ASV compositions. See Fig. S1 for color profiles of bacterial families. Reproduced from the data of a previous study [19]. **B** Taxonomic compositions inferred with shotgun metagenomic sequencing. At each of the 13 time points through the time-series of the target microcosm, the relative abundance of each MAG was estimated based on the normalized read coverage value (reads per kilobase of genome per million reads mapped). **C** Genome size and GC nucleotide content of the MAGs detected in the target microcosm. See panel **A** for colors and symbols.

By targeting the replicate community mentioned above, we compiled shotgun metagenomic data at 13 time points across the time-series [27]. In total, 32 high-quality (> 80 % completeness and < 5 % contamination) metagenome-assembled genomes (MAGs) belonging to 20 genera (16 families; 12 orders) were detected [27] (Figs. 1B-C and 2; Fig. S2; Table S1). As indicated in the amplicon sequencing analysis [19] (Fig. 1A), drastic shifts from taxon-rich community states to oligopolistic states was observed around Day 20 in the shotgun sequencing analysis (Fig. 1B).

After the drastic community compositional change, the system reached a quasi-stable state represented by the dominance of a *Hydrotalea* (Chitinophagaceae) bacterium (Fig. 1B).

The MAG of the *Hydrotalea* was characterized by relatively low GC content (38 %) and relatively small genome size within the community (ca. 3.1 Mb; Fig. 2A). In contrast, the two bacterial MAGs consistently coexisted with the dominant *Hydrotalea* through the time-series (i.e., *Terracidiphilus* and *Mangrovibacter*) had larger genome size (4.2 and 5.4 Mb, respectively; Fig. 1C), characterized by various genes absent from the *Hydrotalea* genome (Fig. 2; Fig. S3). Specifically, the *Terracidiphilus* MAG showed metabolic pathways/processes for degrading plant-derived biopolymers (e.g., cellulose; Fig. 2), potentially surviving as a primary user of polymer compounds within the plant-derived (oatmeal) medium. Meanwhile, the *Mangrovibacter* MAG had pathways/processes related to starch degradation (e.g., amylase) and vitamin-B_12_ transportation, which were absent from the genomes of *Hydrotalea, Terracidiphilus*, and the other MAG (*Rhizomicrobium*) detected on Day 40-60 (Fig. 2).

**Fig. 2.**
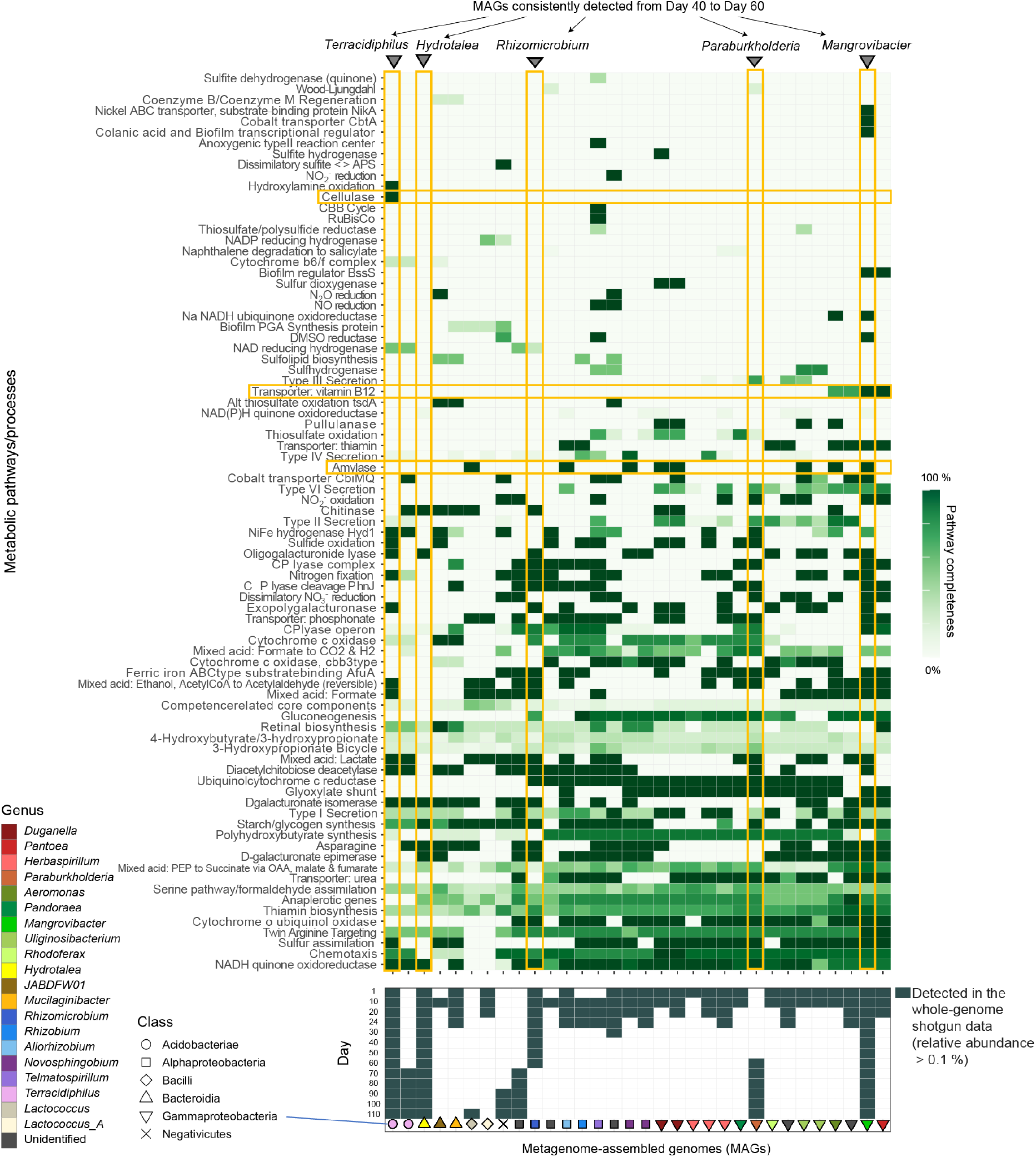
Metabolic pathway/process profiles of the MAGs. KEGG metabolic pathways/profiles of the reconstructed bacterial genomes (MAGs) are shown. The detection (relative abundance > 0.1 %) of each microbial MAG on each day within the shotgun metagenomic data is indicated in the panel below. Only the microbial MAGs with > 80 % completeness and < 5 % contamination were included (Table S1). The five MAGs that co-occurred from Day 40 to 60 and metabolic pathways/processes mentioned in the main text are highlighted. Only the metabolic pathways/processes with highly heterogeneous patterns across microbial MAGs are shown. See Fig. S3 for detailed profiles of the metabolic pathways/processes.

### Multivariate analysis of gene repertoires

Next, we used the shotgun metagenomic data to evaluate how the level of gene repertoire overlap among microbes shifted through time. We anticipated that microbial species with similar resource-use abilities or restrictions have similar genomic structure. Therefore, it is expected that species competing for the same resource tend to form clusters within the space defined based on the principal coordinate analysis (PCoA) of dissimilarity in gene repertoires. For each pair of the 32 MAGs, dissimilarity (Jaccard distance) of gene repertoires was calculated based on the matrix representing the presence/absence of the 6,999 genes annotated with the program Prokka [28]. A PCoA was then performed using the *β*-diversity information (Fig. 3A). At each of the 13 time points, detected MAGs were plotted on the PCoA space.

**Fig. 3.**
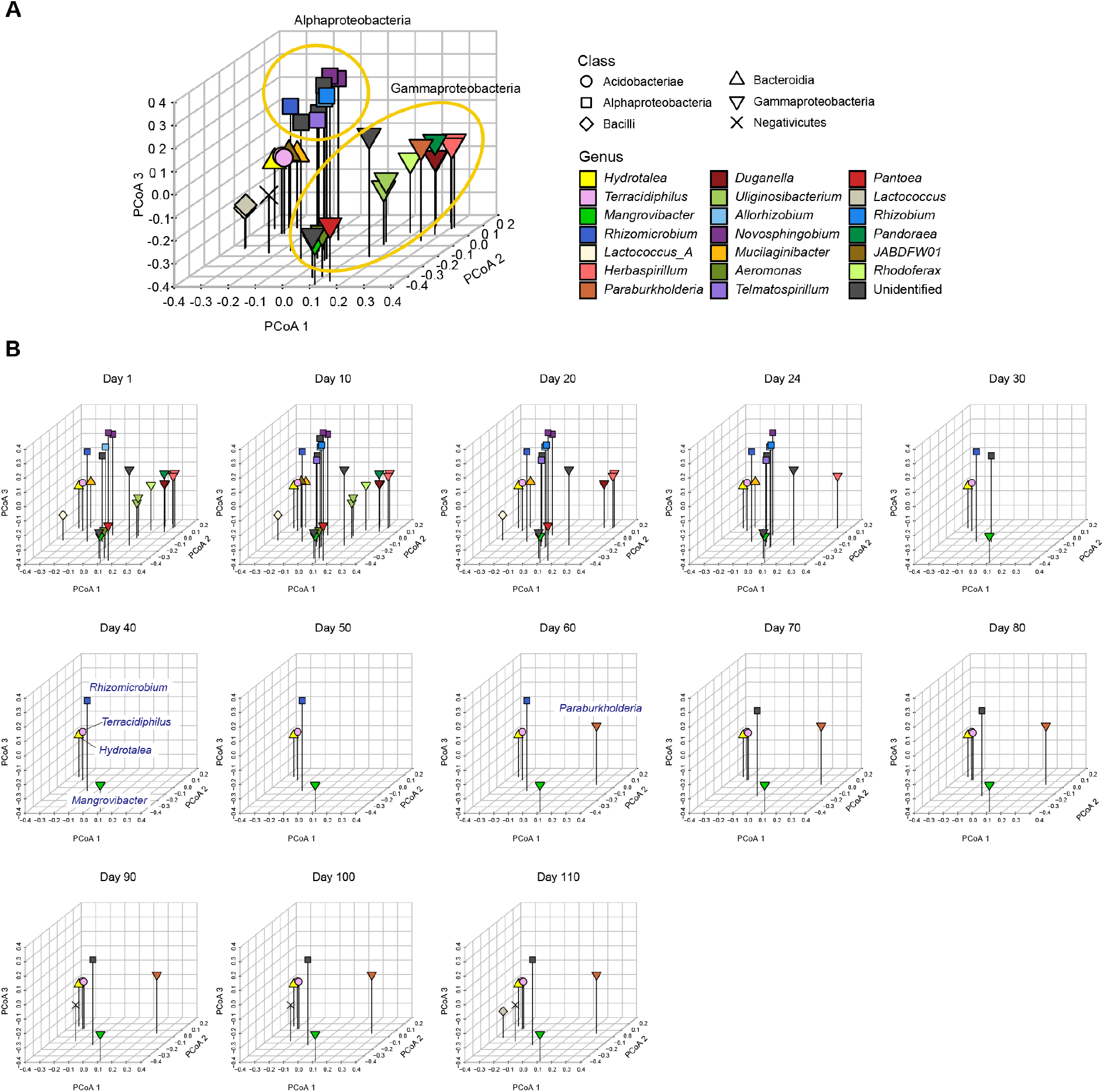
Metagenomic niche space. **A** Distributions of MAGs within metagenomic niche space. Based on dissimilarity in gene repertoires, microbial MAGs that appeared in the time-series of the target microcosm were plotted on the three-dimensional space defined by the principal coordinate analysis (PCoA) of 6,999 genes. **B** Changes in the distributions of microbial MAGs within niche space. At each time point, detected MAGs (relative abundance > 0.1 %) were plotted on the space defined in the multivariate analysis in the in the panel **A**.

Since we did not have *a priori* knowledge of specific metabolic pathways keys to the microbe-to-microbe competition within the experimental microbiome, all datasets were included in this multivariate analysis. Given general characteristics of multivariate analysis based on *β*-diversity metrics, the multivariate reconstruction of ecological niche space depends greatly on the genes whose presence/absence profiles vary among species, while housekeeping genes possessed by most species are expected to contribute little to the multivariate analysis.

We found that alphaproteobacterial and gammaproteobacterial MAGs respectively constituted some clusters within the niche space reconstructed based on the multivariate analysis early in the microbiome dynamics (Day 1-20; Fig. 3B). This state with high niche overlap and potential within-guild competition for resources then collapsed into a simpler community state represented by *Hydrotalea, Mangrovibacter, Terracidiphilus*, and *Rhizomicrobium* as detailed above (Fig. 3B). The space once occupied by many alphaproteobacterial and gammaproteobacterial MAGs remained unoccupied or sparsely occupied after the community compositional collapse. Even when the number of MAGs detectable with our shotgun-metagenomic sequencing increased again late in the time-series, dense aggregations of microbes with similar genomic compositions remained unobserved (Fig. 3B).

### Metagenomic niche overlap

We next quantitatively evaluated dynamics in the magnitude of community-scale niche overlap within the multivariate space (Fig. 3). We developed two types of simple indices for evaluating community-scale niche overlap. The one is defined as the overall mean of gene-repertoire similarity between pairs of MAGs within a community. For a time point, the niche overlap index is calculated as:

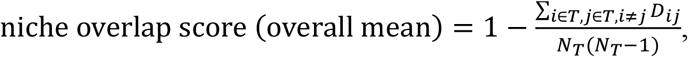

where *T* is the set of MAGs detected on a focal day (relative abundance > 0.1 %), *D*_*ij*_ is the Jaccard metric of dissimilarity [29] in gene compositions, and *N*_*T*_ is the number of MAGs detected on the day. By definition, this niche overlap value based on Jaccard dissimilarity varies from 0 (completely different repertoires of genes in all pairs of MAGs) and 1 (completely identical gene repertoires in all pairs of MAGs), allowing us to evaluate niche overlap levels of target communities within the standardized ranges. The other index is defined as mean value of gene-repertoire similarity with nearest neighbors. The alternative is calculated as:

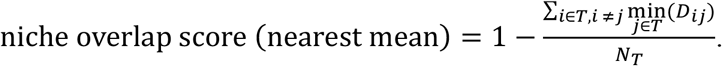

This index can be modified by incorporating the information of the relative abundance of MAGs (*p*_*i*_) as follows:

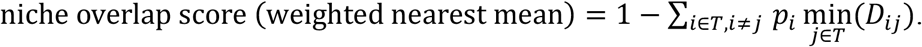

The results indicated that the level of niche overlap was the highest on Day 1 or Day 10 and that it decreased until Day 30 (Fig. 4A). Although the niche overlap score remained low between Day 40 and 60, it increased again late in the microbiome time-series (Fig. 4B). Note that *α*-diversity of the community showed similar temporal shifts and it was significantly associated with the niche overlap indices (Fig. 4B-C). Through the time-series, the estimated niche overlap level was significantly associated with the magnitude of the observed community compositional changes (Fig. 5)

**Fig. 4.**
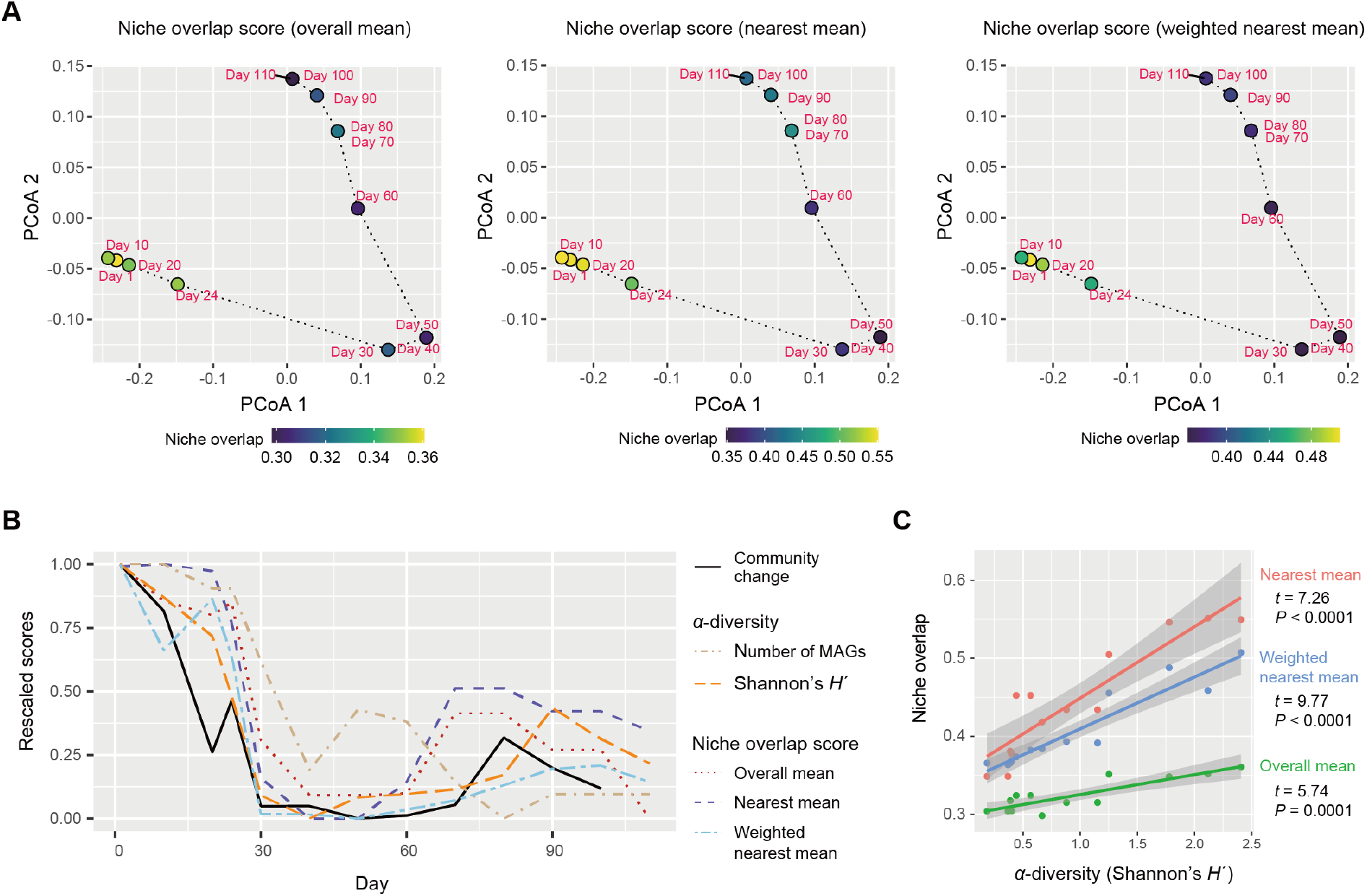
Dynamics of niche-overlap level. **A** Community-level profiles of metabolic pathways/processes and niche overlap index. The niche overlap indices were defined based on the Jaccard similarity/dissimilarity of gene compositions between pairs of the microbial MAGs detected at a target time point. Three types of niche overlap indices are shown on a PCoA surface representing community-level compositions of genes. On the PCoA surface, time points are distributed based on the sum of the gene repertoires of the detected MAGs. **B** Dynamics of niche-overlap levels. Niche overlap scores are shown across the time-series. The magnitude of community compositional changes (Bray-Curtis *β*-diversity between present and next time points through the time-series of the shotgun metagenomic data) and *α*-diversity indices of the communities are shown as well. **C** Relationship between *α*-diversity and niche overlap scores. The lines represent linear regressions (with 95 % confidence intervals).

**Fig. 5.**
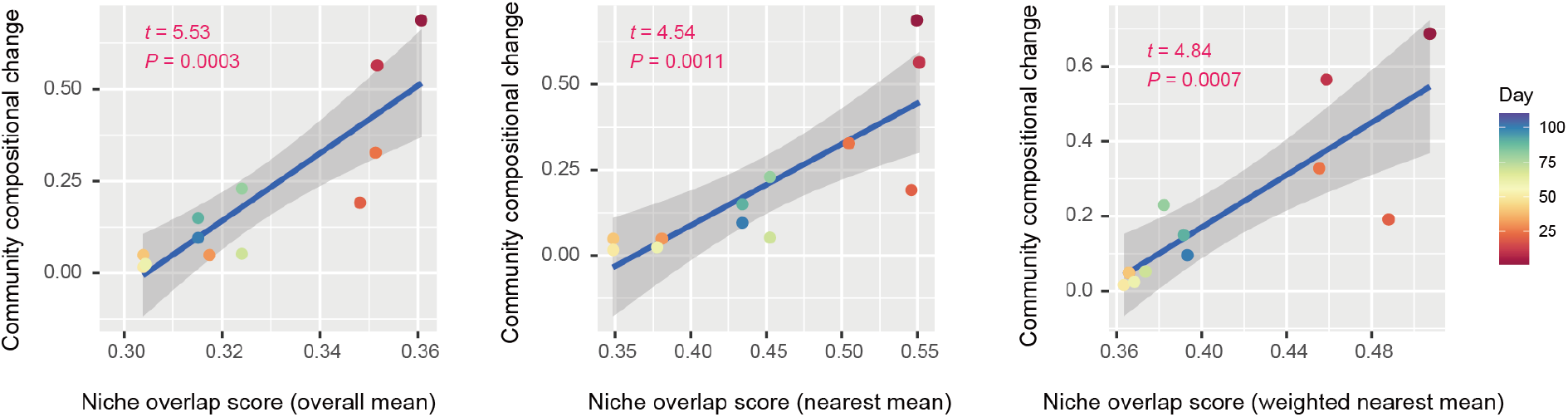
Niche overlap level and community compositional shifts. The magnitude of community compositional changes observed in the microbiome was regressed on each niche overlap index obtained based on the shotgun metagenomic analysis. Niche overlap index at each time point and time-series shifts in community structure (Bray-Curtis *β*-diversity between present and next time points through the time-series of the shotgun metagenomic data) are shown along horizontal and vertical axes, respectively. The regression lines are shown with 95 % confidence intervals.

## DISCUSSION

By developing simple metrics of among-species overlap of gene repertoires, we examined potential relationship between community-scale niche overlap and drastic changes in community structure. Early in the experimental microbiome dynamics, alphaproteobacterial and gammaproteobacterial species were present, resulting in relatively high niche-overlap scores at the community level (Figs. 3 and 4). The quasi-equilibrium state of microbial compositions then collapsed into another quasi-equilibrium represented by a small number of bacteria varying in genome size and metabolic capabilities. Throughout the time-series, higher niche overlap levels entailed greater changes in microbial community compositions (Fig. 5). These findings lead to the working hypothesis that collapse of microbiome structure is predicted by the level of potential niche overlap within multivariate metagenomic space. In light of the “limiting similarity” rule of ecological niches [30], microbial species that exceed a critical limit of genome compositional similarity are expected to compete for the same resources intensively, eventually driving competitive exclusion processes. Thus, as examined in this study, similarity/dissimilarity in genetically determined resource-use properties (i.e., fundamental niches) sets baselines for consequences of interspecific interactions.

The results also indicated that niche overlap level does not necessarily show monotonic decrease through microbial community processes. Although gene-repertoire overlap level and detectable species richness sharply declined early in the microbiome dynamics, both variables gradually increased again around Day 80 (Figs. 1A and 4B). In the resurgence process, however, the dense clusters of alphaproteobacterial or gammaproteobacterial species detected until Day 20 did not appear again within the niche space (Fig. 3B). These observations suggest that once collapsed, microbial communities may not return to previous states with highest levels of niche overlap, but refilling of poorly-used niches can occur under the constraint of limiting similarity within niche space. Although these insights are useful, our present analysis is based only on 13 time points of a microbiome experiment. Due to the limitation, it remained difficult to separate effects of *α*-diversity from those of gene-repertoire overlap (Fig. 4B-C). Thus, the statistical analysis proposed in this study need to be expanded by reducing the cost of metagenomic sequencing as well as by developing more efficient pipelines for the computationally intensive analyses of metagenomic datasets.

The approach of systematically evaluating potential overlap of ecological niches have been previously explored in “community phylogenetics”, in which phylogenetic overdispersion/clustering is evaluated based on null model analysis of random assembly from species pools [5, 31, 32]. In those studies based on phylogenetic analyses, similarity of niches has been inferred based on the assumption that phylogenetically similar species have similar ecological properties (e.g., resource requirements). Nonetheless, given that convergent evolution of ecologically important traits is ubiquitous in the history of life [33–35], the assumption of phylogenetic niche conservatism is not always met [36]. Therefore, because gene repertoires are more direct proxies of species traits than phylogeny, metagenome-based analyses will deepen our understanding of community processes driven by competitive exclusion. Meanwhile, in the present analyses of gene repertoire overlap, we included whole metagenomic datasets of the examined microbes due to the lack of *a priori* insights into the metabolic pathways/processes playing essential roles in interspecific competition for resources. In this respect, our analysis is a preliminary conceptual step for evaluating potential overlap of fundamental niches at the community level. In future studies, analyses excluding housekeeping genes [37, 38] or those focusing on specific functional groups of genes (e.g., carbohydrate degrading genes [39]) may provide more reliable inference of niche overlap.

Because such selection of genes can critically influence threshold niche-overlap values for anticipating abrupt community compositional changes, setting a commonly applicable criterion of choosing target gene sets will help us perform comparative analyses across a wide range of microbial communities.

The simple framework for evaluating overlap of fundamental niches is applicable to diverse types of microbiomes. Given that our Jaccard-dissimilarity-based indices are standardized within the range from 0 to 1, the next crucial step is to examine how threshold niche overlap values for anticipating microbial community collapse vary among different types of ecosystems. Such threshold values can vary among ecosystems depending on their basic levels of sustainable functional redundancy. In our laboratory microbiome, for example, the lack of spatial structure (e.g., refuges for inferior species) and environmental fluctuations (e.g., temperature fluctuations) might have severely limited coexistence of functionally similar species (species with similar metabolic capabilities). In contrast, in human gut microbiomes, spatial complexity [40, 41] and temporally fluctuating environmental conditions [22] may reduce the risk of competitive exclusion, allowing higher levels of niche overlap within communities. Thus, extension of time-series metagenomic analyses to diverse types of ecosystems [42–45] will enhance our knowledge of relationship among ecosystem properties, functional redundancy, and community stability.

While genomic information provides an ultimate platform for inferring fundamental niches [12–14], overlap of gene repertoires may not always result in competitive exclusion of species within communities. Even in a pair of species with similar gene repertoires, differentiation in gene expression patterns may occur to avoid overlap of resource-use patterns between species, allowing coexistence of the two species in an environment. Such differentiation of “realized niches [10]” through phenotypic plasticity is potentially evaluated by transcriptomic or metabolomic analyses [46, 47]. Consequently, integration of (meta)transcriptome and (meta)metabolome analyses [48–50] with metagenome-based analyses will reorganize our understanding of deterministic processes in microbiome dynamics.

## MATERIALS AND METHODS

### Time-series data of experimental microbiomes

We used the experimental system of the microbiome time-series monitoring described in a previous study [19]. In the experiment, microbiomes differing in the magnitude of community compositional shifts were constructed across the six treatments defined by the combinations of two inoculum sources and three types of media. One of the source microbiomes derived from the soil collected from the A layer (0-10 cm in depth) in the research forest of Center for Ecological Research, Kyoto University, Otsu, Japan (34.972 ºN; 135.958 ºE). The other source inoculum was prepared by collecting water from a pond (“Shoubuike”) near Center for Ecological Research (34.974 ºN, 135.966 ºE). Each of the source inocula was introduced into oatmeal (Medium-A), oatmeal-peptone (Medium-B), or peptone (Medium-C) broth media with eight replicates. Thus, in total, 48 experimental microcosms (two source microbiomes × three media × eight replicates) were constructed in a deep-well plate (1000-μL-scale culture in each well). The plate was kept shaken at 1,000 rpm at 23 ºC. After five-day pre-incubation, 200 μL out of the 1,000-μL culture medium was sampled from each well every 24 hours for 110 days. In each sampling event, 200 μL of fresh medium was added to each well so that the total culture volume was kept constant. In total, 5,280 samples (48 communities/day × 110 days) were collected through the time-series experiment. After DNA extraction, the samples were subjected to the amplicon sequencing analysis of the 16S rRNA region [19]. To quantify the speed and magnitude of community shifts through time, the “abruptness” index was calculated through the time-series of each replicate microcosm in each experimental treatment [19]. Specifically, an estimate of the abruptness index for time point *t* was obtained as the Bray-Curtis *β*-diversity between average community compositions from time points *t* – 4 to *t* and those from *t* + 1 to *t* + 5 (i.e., dissimilarity between 5-day time-windows). The Bray-Curtis *β*-diversity [51] was calculated as 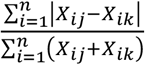, where *X*_*ij*_ and *X*_*ik*_ denoted relative abundance of microbial amplicon sequence variant (ASV) *i* in the compared time windows (*j*, from *t* – 4 to *t*; *k*, from *t* + 1 to *t* + 5). An abruptness score larger than 0.5 indicates that turnover of more than 50 % of community compositions occurred between the time-windows [19].

### Shotgun metagenomics

Focusing on a replicate microcosm in which the most rapid and substantial turnover of community compositions was observed (replicate no. 5 of Water/Medium-A treatment; Fig. S1), shotgun metagenomic sequencing was conducted by targeting 13 samples (Day 1, 10, 20, 24, 30, 40, 50, 60, 70, 80, 90, 100, and 110) as described elsewhere [27]. Specifically, each DNA sample was processed with Nextera XT DNA Library Preparation Kit (Illumina) and sequenced with the DNBSEQ-G400 (BGI; 200-bp paired-end sequencing). From the output data, sequencing adaptors were removed using Cutadapt [52] 2.5 and quality filtering was performed with Fastp [53] 0.21.0: ca. 10 Gb/sample was subjected to the analysis [in total, 159.96 Gb (1000.301 M reads)]. The sequences of each sample were assembled with metaSPAdes [54] 3.15.2. Binning was then performed with MetaWRAP [55] 1.3.2, followed by quality assessing with CheckM [56] 1.1.3. The identity between MAGs were calculated using FastANI [57] 1.33 and MAGs with > 99 % identity were dereplicated through the time-series (Table S1). In the dereplication, the MAGs with the highest completeness and N50 statistics were selected as representative MAGs. Read-coverage was then calculated with CoverM [58] 0.6.0, followed by taxonomic annotation was performed using GTDB-Tk [59, 60] 1.6. Only the MAGs with > 80 % completeness and < 5 % contamination were used in the downstream analyses. Gene annotation was performed with Prokka [28] 1.14.6, yielding 6,999 annotated genes (Data S1). To conduct additional functional annotation of genes, the orthology numbers of Kyoto Encyclopedia of Genomes (KEGG) were retrieved using GhostKOALA [61] 2.2. For respective microbial MAGs (bins), completeness of metabolic pathways was estimated with KEGG decoder [62] 1.3. Based on the matrix representing KEGG metabolic pathway/process profiles of respective MAGs (Data S2), a heatmap showing pathway/process completeness was drawn (Fig. S3).

### Background environmental conditions

For the 13 samples subjected to the shotgun metagenomic analysis, concentrations of ammonium (NH_4_^+^) and nitrate (NO_3_^−^) were measured to obtain supplementary information of background environmental conditions. Colorimetric methods with a modified indophenol reaction [63, 64] and the VCl3/Griess assay were applied for the measurements of NH_4_^+^ and NO_3_^−^, respectively. Samples were run in triplicates via a standard addition method to account for individual matrix effects [65].

### Multivariate analysis of the metagenomic space

Based on the whole matrix representing the profiles of the 6,999 genes (Data S1), the Jaccard metric of distance was calculated for each pair of the 32 microbial MAGs (*D*_*ij*_, where *i* and *j* represent MAGs). The Jaccard-distance estimates were then used to perform a principal coordinate analysis (PCoA). Using the obtained principal coordinate scores, all the microbial MAGs detected through the time-series were plotted on a multivariate space consisting of the first three PCoA axes (PCoA 1, PCoA 2, and PCoA 3). At each time point, the MAGs detected with the shotgun metagenomic sequencing (defined as the MAGs whose relative abundance is greater than 0.1 %) was plotted on the three-dimensional space defined with the PCoA axes.

### Evaluation of niche overlap level

The community-scale magnitude of potential niche overlap among species was evaluated based on the shogun metagenomic sequencing dataset. We developed two types of simple indices for evaluating community-scale niche overlap as detailed in the Results section. To test whether a high level of fundamental-niche overlap is observed prior to drastic changes in microbial community structure, we examined relationship between the above niche overlap index and time-series shifts in community structure (Bray-Curtis *β*-diversity between present and next time points through the time-series of the shotgun metagenomic data).

## ACKNOWLEDGEMENTS

Computation time was provided by the SuperComputer System, Institute for Chemical Research, Kyoto University. This work was financially supported by JST PRESTO (JPMJPR16Q6), Human Frontier Science Program (RGP0029/2019), JSPS Grant-in-Aid for Scientific Research (20K20586), NEDO Moonshot Research and Development Program (JPNP18016), and JST FOREST (JPMJFR2048) to H.T., JSPS Grant-in-Aid for Scientific Research (20K06820 and 20H03010) to K.S., and JSPS Fellowship to H.F. and A.C.

## AUTHOR CONTRIBUTIONS

H.T. designed the work with H.F. H.F. and A.C. performed experiments. H.F. analyzed the data with Y.O., and H.T. H.F. and H.T. wrote the paper with all the authors.

## DATA AVAILABILITY STATEMENT

The 16S rRNA sequencing data reported in a previous study [19] are available from the DNA Data Bank of Japan (DDBJ) with the accession number DRA013352, DRA013353, DRA013354, DRA013355, DRA013356, DRA013368 and DRA013379. The shotgun metagenomic data reported previous [27] are available with the DDBJ accession number DRA013382. The microbial community data are deposited at our GitHub repository (https://github.com/hiroakif93/MTS_nicheSpace) [to be publicly available after acceptance of the paper]. The matrices of the shotgun metagenomic data are available as Data S1 and 2. All the scripts used to analyze the data are available at the GitHub repository (https://github.com/hiroakif93/MTS_nicheSpace) [to be publicly available after acceptance of the paper].

