## Supplementary Figures for "Metagenomic analysis of ecological niche overlap and community collapse in microbiome dynamics"

**This PDF file includes:**

Supplementary Figures S1-3

Supplementary Table S1

**Supplementary Information included in a separate file:**

Supplementary Data S1-2

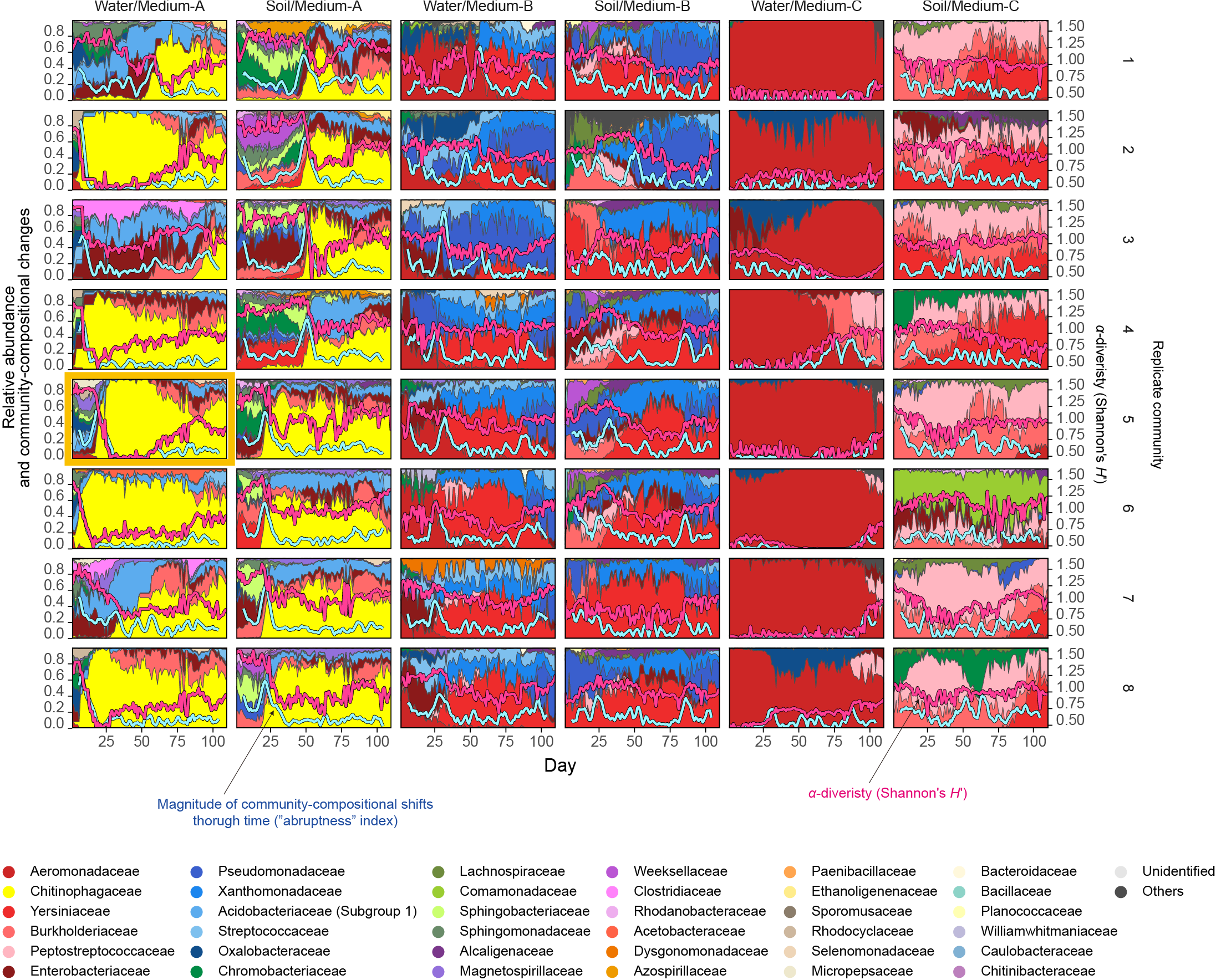

**Fig. S1 | Dynamics of family-level community structure**. The dynamics of microbial family-level compositions were visualized based on the 16S rRNA sequencing data of the previous study [1]. The replicate microcosm (replicate no. 5 in Water/Medium-A treatment), which is subjected to the shotgun metagenomic sequencing analysis, is highlighted. The blue line represents the speed and magnitude of community compositional changes around each time point (“abruptness” index [1]; see Methods). The red line indicates *α*-diversity (Shannon’s *H’*) of microbial ASVs [1]. Reproduced from the data of a previous study [1].

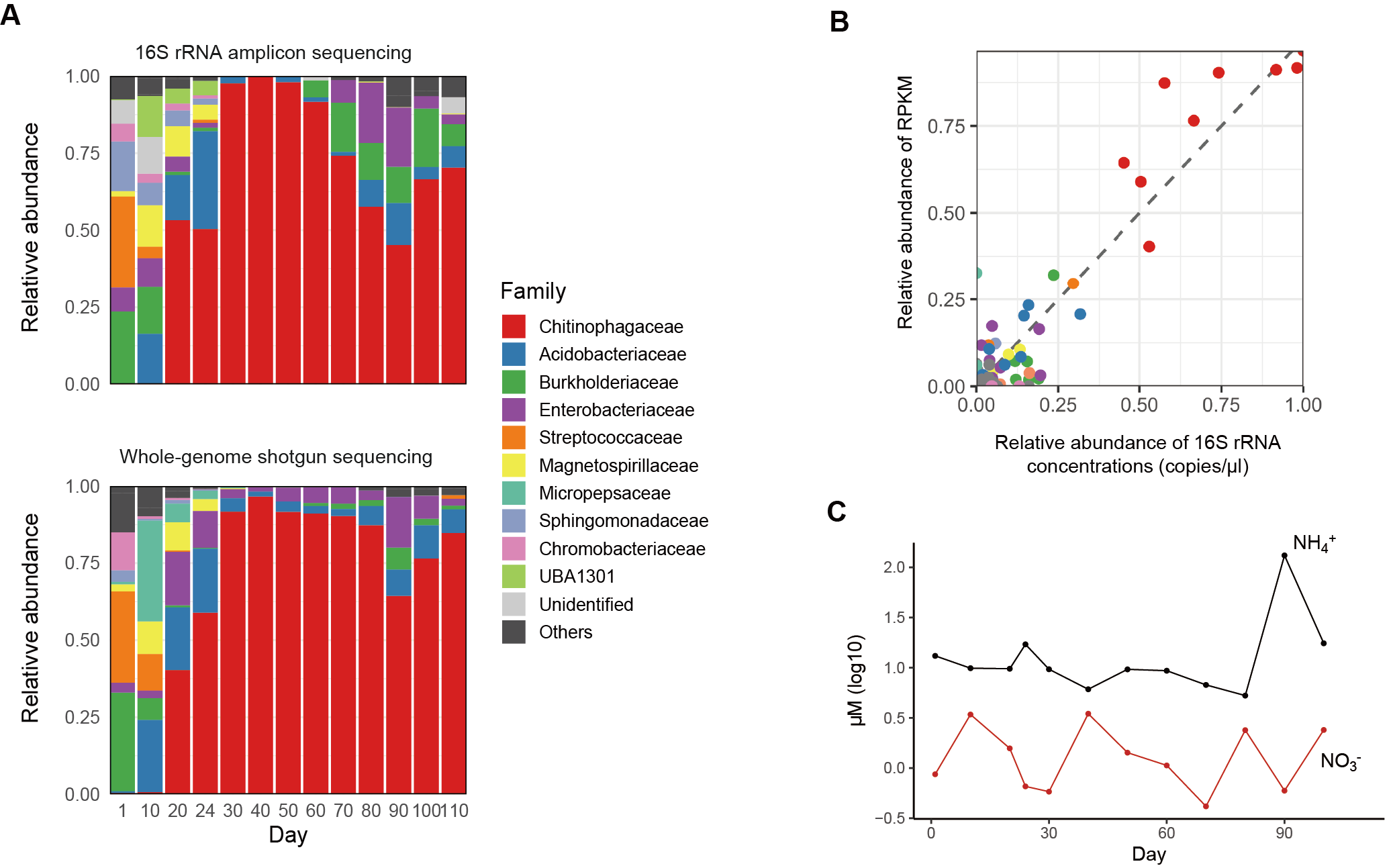

**Fig. S2 | Overview of the shotgun metagenomic sequencing data.** **A** Comparison of relative abundance of bacterial taxa (families) between 16S rRNA amplicon sequencing [1] (reproduced from the previous study [1]; top) and shotgun metagenoimc sequencing (reproduced from the previous study [2]; bottom). **B** Correlation between the family-level relative abundance of 16S rRNA and shotgun metagenomic sequencing data (Spearman’s correlation; *ρ* = 0.667, df = 794, *P* < 0.05). Each point represents each family at each time point. **C** Background chemical properties. Changes in NO_3_^−^ and NH_4_^+^ concentrations in the ecosystem are shown for the time points with shotgun metagenomic metagenomic data.

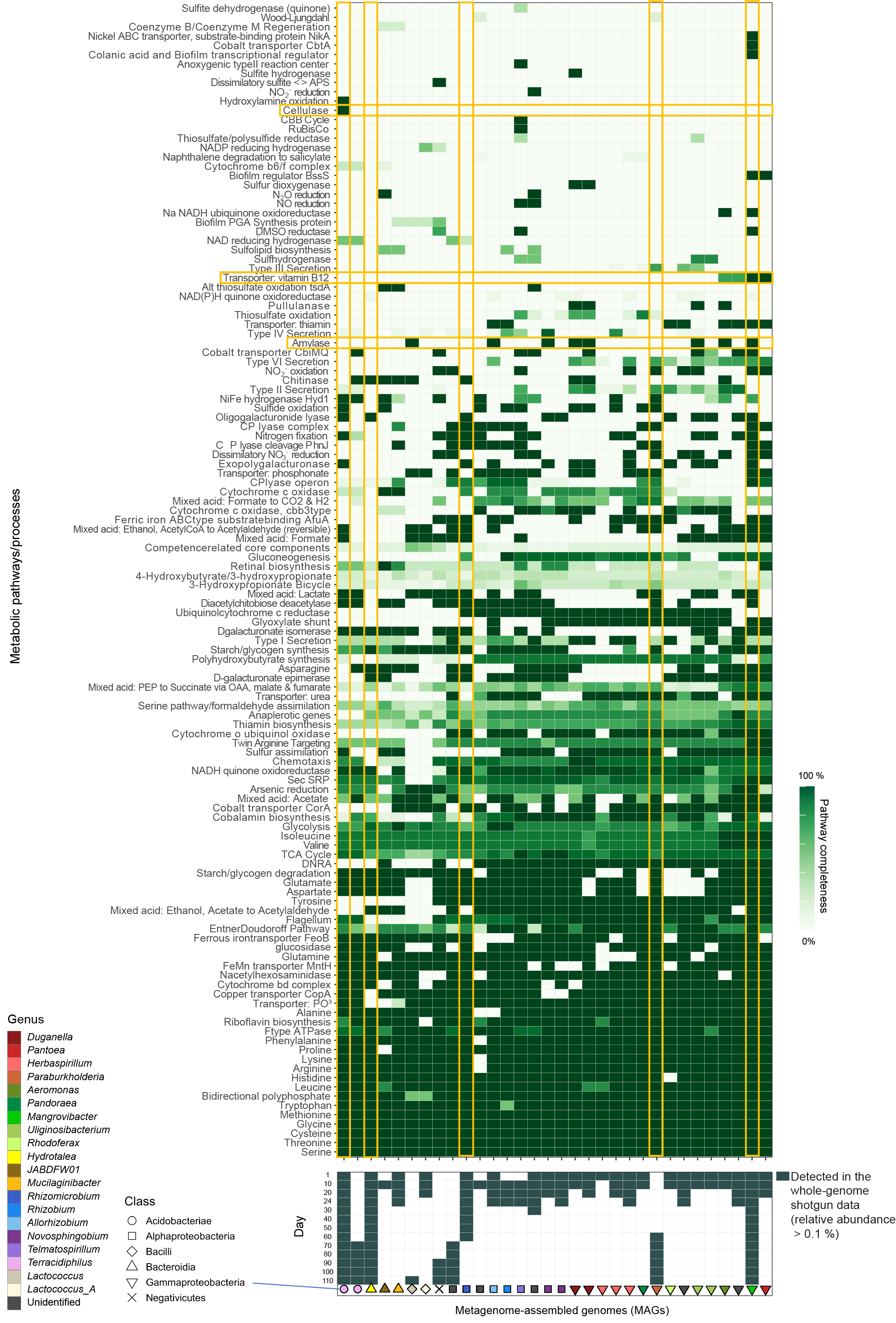

**Fig. S3 | Detailed information of the metabolic pathway/process profiles of the MAGs.** The KEGG metabolic pathways/processes of the reconstructed bacterial genomes (MAGs) are shown. The detection (relative abundance > 0.1 %) of each microbial MAG on each day within the shotgun metagenomic data is indicated in the panel below. Only the microbial MAGs with > 80 % completeness and < 5 % contamination were included (Supplementary Table 1). The five MAGs that co-occurred from Day 40 to 60 and metabolic pathways/profiles mentioned in the main text are highlighted. The detailed definition of the KEGG metabolic pathways/processes is available at https://github.com/bjtully/BioData/blob/master/KEGGDecoder/KOALA_definitions.txt.

**Table S1 | List of microbial MAGs detected in the whole-genome shotgun metagenomics.** For each representative MAGs, estimates of completeness and contamination rates of the genome reconstruction as well as estimated genome size, GC nucleotide contents, coding density, and number of predicted genes are shown.

| MAG ID | Phylum | Class | Order | Family | Genus | Species | Completeness (%) | Contamination (%) | Genome size (bp) | GC content (%) | Coding density (%) | Number of predicted genes |
| --- | --- | --- | --- | --- | --- | --- | --- | --- | --- | --- | --- | --- |
| MTS-010_bin.016 | Proteobacteria | Gammaproteobacteria | Burkholderiales | Burkholderiaceae | *Duganella* |  | 99 | 0.33 | 5548277 | 62.5 | 91.6 | 4993 |
| MTS-001_bin.010 | Proteobacteria | Alphaproteobacteria | Sphingomonadales | Sphingomonadaceae | *Novosphingobium* |  | 99.09 | 3.29 | 5636625 | 64.3 | 90.31 | 4976 |
| MTS-001_bin.011 | Proteobacteria | Alphaproteobacteria | Rhodospirillales | Magnetospirillaceae |  |  | 99.38 | 0 | 3951389 | 62.6 | 91.44 | 3661 |
| MTS-010_bin.001 | Proteobacteria | Gammaproteobacteria | Burkholderiales | Chromobacteriaceae |  |  | 99.57 | 0.85 | 4573844 | 52.1 | 85.31 | 4255 |
| MTS-001_bin.013 | Proteobacteria | Alphaproteobacteria | Sphingomonadales | Sphingomonadaceae | *Novosphingobium* |  | 98.86 | 0.69 | 4198163 | 62.4 | 89.89 | 3913 |
| MTS-010_bin.008 | Firmicutes | Bacilli | Lactobacillales | Streptococcaceae | *Lactococcus_A* | *Lactococcus_A raffinolactis* | 98.11 | 1.54 | 2372475 | 39.6 | 87.86 | 2327 |
| MTS-040_bin.003 | Proteobacteria | Gammaproteobacteria | Enterobacterales | Enterobacteriaceae | *Mangrovibacter* |  | 99.56 | 0.27 | 5404296 | 52.2 | 87.61 | 5182 |
| MTS-001_bin.016 | Proteobacteria | Gammaproteobacteria | Burkholderiales | Burkholderiaceae | *Duganella* | *Duganella* sp003217375 | 99.92 | 1.19 | 6479266 | 62.6 | 92.81 | 5830 |
| MTS-090_bin.001 | Bacteroidota | Bacteroidia | Chitinophagales | Chitinophagaceae | *Hydrotalea* |  | 98.52 | 0 | 3112189 | 38 | 85.06 | 2521 |
| MTS-001_bin.018 | Proteobacteria | Gammaproteobacteria | Burkholderiales | Burkholderiaceae | *Herbaspirillum* |  | 84.16 | 0.74 | 3632523 | 54.3 | 89.11 | 3567 |
| MTS-001_bin.019 | Proteobacteria | Gammaproteobacteria | Burkholderiales | Burkholderiaceae | *Rhodoferax* |  | 96.25 | 1.12 | 3077095 | 57 | 93.48 | 3084 |
| MTS-010_bin.009 | Proteobacteria | Gammaproteobacteria | Burkholderiales | Rhodocyclaceae | *Uliginosibacterium* |  | 99.38 | 0.95 | 5092993 | 57 | 89.37 | 4744 |
| MTS-001_bin.020 | Proteobacteria | Gammaproteobacteria | Enterobacterales | Aeromonadaceae | *Aeromonas* | *Aeromonas hydrophila* | 98.88 | 0.29 | 5006854 | 61.1 | 87.34 | 4561 |
| MTS-001_bin.021 | Proteobacteria | Gammaproteobacteria | Burkholderiales | Rhodocyclaceae | *Uliginosibacterium* |  | 87.71 | 3.73 | 4570696 | 58.4 | 87.36 | 4787 |
| MTS-020_bin.006 | Proteobacteria | Gammaproteobacteria | Enterobacterales | Aeromonadaceae |  |  | 100 | 0 | 3907279 | 49.1 | 90.11 | 3725 |
| MTS-001_bin.004 | Bacteroidota | Bacteroidia | Sphingobacteriales | Sphingobacteriaceae | *Mucilaginibacter* | *Mucilaginibacter oryzae* | 97.62 | 0.4 | 5891481 | 44.3 | 87.99 | 5074 |
| MTS-001_bin.005 | Proteobacteria | Gammaproteobacteria | Enterobacterales | Enterobacteriaceae | *Pantoea* | *Pantoea* sp011752625 | 100 | 1.55 | 6025599 | 52.8 | 88.07 | 5560 |
| MTS-001_bin.006 | Proteobacteria | Gammaproteobacteria | Burkholderiales | Burkholderiaceae | *Herbaspirillum* |  | 99.82 | 1.29 | 6377155 | 56.4 | 88.72 | 5610 |
| MTS-001_bin.007 | Proteobacteria | Gammaproteobacteria | Burkholderiales | Burkholderiaceae | *Herbaspirillum* |  | 99.95 | 0.6 | 5673370 | 62.1 | 88.02 | 5070 |
| MTS-010_bin.015 | Proteobacteria | Alphaproteobacteria | Rhizobiales | Rhizobiaceae | *Allorhizobium* |  | 98.31 | 0.84 | 5665159 | 55.1 | 88.93 | 5363 |
| MTS-001_bin.009 | Proteobacteria | Gammaproteobacteria | Burkholderiales | Burkholderiaceae | *Pandoraea* |  | 99.55 | 0.96 | 6402802 | 63.5 | 88.08 | 5703 |
| MTS-010_bin.010 | Proteobacteria | Alphaproteobacteria | Rhizobiales | Phreatobacteraceae |  |  | 82.3 | 0.98 | 3278745 | 59.7 | 90.8 | 3602 |
| MTS-010_bin.011 | Proteobacteria | Alphaproteobacteria | Rhizobiales | Rhizobiaceae | *Rhizobium* |  | 96.14 | 1.99 | 6292370 | 60.5 | 87.24 | 6284 |
| MTS-010_bin.012 | Proteobacteria | Alphaproteobacteria | Micropepsales | Micropepsaceae | *Rhizomicrobium* |  | 98.9 | 0.74 | 3592561 | 58.6 | 88.84 | 3388 |
| MTS-020_bin.011 | Proteobacteria | Alphaproteobacteria | Rhodospirillales | Magnetospirillaceae | *Telmatospirillum* |  | 99.5 | 1.49 | 5604519 | 61.3 | 89.37 | 5178 |
| MTS-010_bin.005 | Bacteroidota | Bacteroidia | Chitinophagales | Chitinophagaceae | JABDFW01 |  | 88.53 | 0.25 | 4844654 | 44 | 87.52 | 4047 |
| MTS-060_bin.002 | Acidobacteriota | Acidobacteriae | Acidobacteriales | Acidobacteriaceae | *Terracidiphilus* |  | 97.41 | 0.86 | 4157144 | 60.1 | 87.38 | 3415 |
| MTS-090_bin.004 | Proteobacteria | Gammaproteobacteria | Burkholderiales | Burkholderiaceae | *Paraburkholderia* | *Paraburkholderia tropica* | 99.6 | 2.1 | 8236275 | 65 | 86.47 | 7367 |
| MTS-100_bin.002 | Acidobacteriota | Acidobacteriae | Acidobacteriales | Acidobacteriaceae | *Terracidiphilus* |  | 99.14 | 0.86 | 4174131 | 58.4 | 87.74 | 3312 |
| MTS-090_bin.002 | Proteobacteria | Alphaproteobacteria | Acetobacterales | Acetobacteraceae |  |  | 99.5 | 0.5 | 4089046 | 68.2 | 88.35 | 3679 |
| MTS-090_bin.007 | Firmicutes_C | Negativicutes | Selenomonadales | Selenomonadaceae |  |  | 99.91 | 0.63 | 3554726 | 41.6 | 88.47 | 3344 |
| MTS-110_bin.003 | Firmicutes | Bacilli | Lactobacillales | Streptococcaceae | *Lactococcus* | *Lactococcus lactis* | 99.62 | 0.51 | 2257933 | 34.9 | 87.27 | 2228 |
